# Lhx2 regulates temporal changes in chromatin accessibility and transcription factor binding in retinal progenitor cells

**DOI:** 10.1101/238279

**Authors:** Cristina Zibetti, Sheng Liu, Jun Wan, Jiang Qian, Seth Blackshaw

**Affiliations:** Solomon H. Snyder Department of Neuroscience, Johns Hopkins University School of Medicine, Baltimore, MD, 21205, USA; Department of Ophthalmology, Johns Hopkins University School of Medicine, Baltimore, MD, 21205, USA; Department of Neurology, Johns Hopkins University School of Medicine, Baltimore, MD, 21205, USA; Center for Human Systems Biology, Johns Hopkins University School of Medicine, Baltimore, MD, 21205, USA; Institute for Cell Engineering, Johns Hopkins University School of Medicine, Baltimore, MD, 21205, USA; Department of Medical and Molecular Genetics, Indiana University School of Medicine, Indianapolis IN 46202

## Abstract

Retinal progenitor cells (RPCs) pass through multiple stages of developmental competence, where they successively acquire and lose the ability to generate individual cell subtypes. To identify the transcriptional regulatory networks that control these transitions, we conducted epigenomic and transcriptomic profiling of early and late-stage RPCs and observed a developmentally dynamic landscape of chromatin accessibility. Open chromatin regions that showed stage-specificity, as well as those shared by early and late-stage RPCs, were selectively targeted by the homeodomain factor Lhx2, which is expressed throughout retinal neurogenesis but also regulates many stage-specific processes in RPCs. Stage-specific Lhx2 binding sites were frequently associated with target sites for transcription factors that are preferentially expressed in either early or late-stage RPCs, and which were predicted to possess pioneer activity. *Lhx2* loss of function in RPCs led to a loss of chromatin accessibility at both direct Lhx2 target sites and more broadly across the genome, as well as a loss of binding by transcription factors associated with stage-specific Lhx2 target sites. These findings demonstrate a central role for Lhx2 in control of chromatin accessibility in RPCs, and identify transcription factors that may guide stage-specific target site selection by Lhx2.

**Summary:** Lhx2 is a central regulator of chromatin accessibility in retinal progenitor cells, and interacts with stage-specific transcription factors to regulate genes that are dynamically expressed during retinal neurogenesis.

## Introduction

During the course of neurogenesis, neural progenitors move through different states of developmental competence, in which they successively acquire and lose their ability to generate individual cell subtypes. This process, also known as temporal patterning, occurs throughout the nervous system in both vertebrates and invertebrates (*1, 2*). These transitions are largely cell-autonomous and proceed in a unidirectional and irreversible manner, and are driven by changes in transcription factor (TF) expression. The murine retina is an excellent model for mechanistic analysis of temporal patterning, as the generation of all major cell types is controlled through this process. A limited number of TFs have been identified that show differential expression in early and late-stage retinal progenitor cells (RPCs), and which also control changes in developmental competence *(3–6)*. Other TFs expressed throughout retinal neurogenesis, such as the LIM homeodomain factor *Lhx2*, are nonetheless essential for regulating multiple stages of RPC competence. Lhx2 is required for the transition of early-stage RPCs from an immature state, in which they can generate retinal ganglion cells (RGCs), to a more mature state in which they can no longer do so (*7*). In contrast, in late-stage RPCs, Lhx2 is essential for generation of late-born Muller glia and does this in part by enhancing Notch signaling *(8, 9)*. Lhx2 regulates these temporally dynamic processes in parallel with transcriptional programs common to early and late-stage RPCs, where it maintains proliferative and neurogenic competence, and represses expression of genes that are specific to the anterodorsal hypothalamus and thalamic eminence *(10)*.

The mechanism by which these highly stage-specific TFs interact with broadly expressed factors such as Lhx2 to regulate retinal cell fate specification is unknown, and a broader picture of the transcriptional regulatory networks that control RPC competence is likewise lacking. In this study, in order to identify *cis* and trans-acting factors that regulate temporal patterning in the developing retina, we globally profiled chromatin accessibility and mRNA expression in both early and late-stage RPCs, using ATAC-Seq and RNA-Seq, and identify Lhx2 as a central regulator of changes in chromatin accessibility that occur in RPCs during the course of neurogenesis.

## Results

### Lhx2 binding sites are highly enriched in open chromatin in RPCs

Fluorescence-activated cell sorting (FACS) was used to isolate RPCs from mice expressing the RPC-specific *Chx10-Cre:eGFP* transgene (*11*) from embryonic day (E)14 and postnatal day (P)2 retina, representing early and late stages of RPC competence, respectively (*1*). The great majority of GFP-positive cells expressed the RPC-specific markers Chx10, Ki67 and Ccnd1, while these were absent from the GFP-negative postmitotic fraction (Fig. S1A-G). Flow-sorted cells were profiled by RNA-Seq and ATAC-Seq, relying on direct *in vitro* transposition of sequencing adapters into native chromatin *(12)*. ChlP-Seq was then performed on a select transcription factor candidate, identified in the preliminary screening of the ATAC-Seq derived open chromatin regions. ChlP-Seq binding sites for the candidate factor were ultimately integrated and compared with the age-matched RNA-Seq and ATAC-Seq profiles from control and conditional *Lhx2* knockout retinas (Fig.1 A, S1H, I). ATAC-Seq data was highly reproducible among experimental replicates (Fig. S2A-D), and identified open chromatin regions (OCRs) that were shared between time points, or were specific to either E14 or P2 (Fig. 1B). These represent 2.4%, 0.72%, and 0.80% of the genome, respectively.

We next scanned OCRs common to E14 and P2 RPCs for candidate TF binding sites. Clustering of the assigned position weight matrices (PWM) revealed that homeobox-containing TFs constitute the prevalent cluster, representing 284/373 putative matches (Fig. 1C, highlighted in the inset), followed by C2H2-type, GC-rich and KLF zinc finger proteins; followed by POU-homeodomain factors, MADS box and CCAAT-binding factors. The consensus motif for the insulator protein CTCF, involved in controlling chromatin looping, is also highly enriched in OCRs. To pinpoint the individual TFs that showed strong enrichment in common OCRs, we examined the gene expression level of the TFs in RPCs. Interestingly, only a few of the TFs that were the top predicted matches for the homeobox-containing factors were detectably expressed in RPCs by RNA-Seq (Fig. 1D), and many represented close variants of the Lhx2 consensus sequence (Fig. 1E, S2E). Indeed, among the TFs that are expressed in RPCs, Lhx2 consensus site was identified as the most represented TF motif in OCRs common to early and late RPCs (Fig. 1F, Auxiliary table 1), and variably represented in stage-specific OCRs (Auxiliary table 2,3). Lhx2 occupancy resulted into detectable footprints at both stages (Fig. 1G). Finally, ChIP-Seq was performed for Lhx2 in E14 and P2 mouse retina. Integration with age-matched ATAC-Seq profiles from flow-sorted RPCs demonstrated that Lhx2 ChIP-Seq peaks do indeed overlap with many of OCRs (Fig.1H, 2A, S3A-G).

### Lhx2 target site selection in RPCs is dynamic during the course of retinal neurogenesis

Lhx2 ChIP-Seq peaks are predominantly located in intragenic and intronic regions of the genome (Fig. S4A), are enriched in promoter regions, 5’UTR, exons, and ncRNA at both time points (Fig. 2B), and are enriched for functions related to control of retinal development (Fig. S4B,C). The Lhx2 ChIP-Seq data (Fig. S3G) was then compared to a manually curated list of genes that showed enriched expression in individual retinal cell types, which also included RNA-Seq data from E14 and P2 RPCs (Auxiliary table 4,5). Lhx2 ChIP-Seq peaks were preferentially associated with genes that were selectively expressed in late-stage RPCs, Müller glia, and the embryonic anterodorsal hypothalamus at both E14 and P2 (Fig. 2C, Auxiliary table S5). Peaks found only at E14 were also selectively associated with genes expressed in early-stage RPCs (Fig. 2D), while genes enriched in late-stage RPCs were overrepresented in the P2 ChIP-Seq dataset (Fig. 2D).

To identify genes whose expression is directly regulated by Lhx2 in early and late-stage RPCs, Lhx2 targets identified by ChIP-Seq were compared to the RNA-Seq data from the E14 and P2 GFP-positive fractions and to the expression profiles obtained from Lhx2-deficient cells isolated at E14 (*Chx10-Cre;Lhx2^lox/lox^* retinas) (Fig. 2D,E, Fig. S5A-C) and P2 (*Lhx2^lox/lox^* RPCs electroporated with pCAG-Cre-GFP at P0 and isolated by FACS at P2) (Fig. 2D,F, Fig. S5D-F). In total, more than 30% of the Lhx2 peaks shared between time points and 20% of the stage-specific peaks were associated with differentially expressed genes (DEG) detected from the comparison between E14 and P2 RPCs, potentially accounting for 10 to 14% of the differentially expressed transcriptome.

Most of the genes that were differentially expressed between early and late RPCs, as well as those that were enriched in early and late RPCs compared to age-matched GFP-negative postmitotic cells, were associated with stage-specific Lhx2 peaks (Auxiliary table 4). Importantly, 47% of all genes that were differentially expressed between E14 and P2 RPCs also showed altered expression in *Lhx2* cKO samples, at one or both of these timepoints (Fig. S5G, Auxiliary table 6,7). Examples of such temporally dynamic and Lhx2-regulated genes include *Ndnf*, which is upregulated in P2 *Lhx2* cKO cells, and the late-stage RPC and MG-enriched *Car2* locus, which is downregulated in P2 *Lhx2* cKO cells (Fig. S5H). These results show that Lhx2 plays a central role in controlling temporally dynamic gene expression in RPCs.

### Lhx2 selectively targets OCRs and regulates genes associated with RPC function

Lhx2 binding sites are predominantly associated with open chromatin. In E14 RPCs, 4.3% of OCRs overlapped with Lhx2 ChIP-Seq peaks and 3.5% of P2 OCRs were associated with age-matched Lhx2 peaks, with a higher fraction of closely co-localizing peaks at E14 compared to P2 (Auxiliary table 6). The majority of Lhx2 target sites fall in OCRs: 76.7% of the E14 Lhx2 cis-regulatory sites fall in OCRs in E14 GFP-positive RPCs, while at P2 61.7% of the P2 ChIP-Seq peaks co-occur with P2 GFP-positive OCRs (Fig. S4D, Auxiliary table 7). Notably, Lhx2 ChIP-Seq peaks were preferentially co-localized with positioned nucleosomes at E14 (Fig. 2G) but bimodal at P2 (Fig. 2H) suggesting Lhx2 may compete for nucleosome binding in early-stage RPCs (Fig. 2G).

Overall, 9.9% of the nucleosome-interspersed OCRs identified in E14 RPCs, and 9.3% of those identified at P2, are associated with bidirectionally-transcribed Lhx2 binding sites. These represent 19.3% of the high confidence Lhx2 ChIP-Seq peaks at E14 and 17.4% at P2. Lhx2 targets that flanked positioned nucleosomes within OCRs were enriched at 5’UTR and promoter regions at both E14 (Fig. 2G) and P2 (Fig. 2H) and coupled with bimodally distributed, likely active enhancers, marked by H3K27ac peaks at P2. Since exons and ncRNAs were also found among the enriched categories of Lhx2 target sites at both ages, it is likely that a subset of these *cis*-regulatory regions may encode enhancer-associated RNAs (eRNAs).

Lhx2 targets in E14 RPCs were associated with genes involved in regulatory RNA pathways, promoter opening and cell cycle checkpoints (Fig. 2I) and those in P2 RPCs population were enriched for functional categories related cell cycle progression, Notch signaling and DNA replication, as well as phenotypes such as retina hypoplasia (Fig. 2J). Enrichment for these functional categories was confirmed by comparison with *Lhx2*-dependent gene sets identified by RNA-Seq analysis of the age-matched *Lhx2* cKO samples.

### Identification of TF motifs associated with Lhx2 ChIP-Seq peaks

We next investigated which TFs could co-localize with Lhx2 ChIP-Seq peaks at E14 and P2, or at each individual timepoint. The Lhx2 consensus sequence and closely related sequences were over-represented in peaks present at both E14 and P2 (Fig. 3A,B). Multiple motifs were found within a broad range of similarity to the known Lhx2 consensus (Pearson’s correlation > 0.6), possibly underlying differences in affinity and/or combinatorial interaction with other transcription factors (Fig. S6A-I).

In the E14 retina, the SoxB1 related family members (Sox2) *(15)* and MADS transcription factors were found at E14 (Fig. 3A) while Kruppel like factors (Klf9/13) *(13)*, NF-I family members (Nfia/Nfib/Nfic/Nfix) *(16)*, SoxE (Sox8/9) *(17)* and Ascl1 *(18)* were found at P2 (Fig.3B). When only known TF motifs are considered, (Fig. S7A,B), preferential representation of homeobox/bHLH transcription factors was observed at E14 (Fig. 3C) and NF-I, CTCF and Forkhead family members observed at P2 (Fig. 3D).

### Lhx2 broadly regulates chromatin accessibility in RPCs

We then tested whether loss of function of Lhx2 affected chromatin accessibility in RPCs. We first analyzed ATAC-Seq profiles from control samples at E14 (whole retina) and P2 (FACS-isolated cells electroporated at P0) (Fig. S8A-G), and observed high overall correlation with ATAC-Seq profiles from age-matched purified RPCs. By comparing these to ATAC-Seq profiles from E14 and P2 *Lhx2* cKO samples, we observed an extensive reduction in OCRs (Fig. S8F,G).

As expected, we observed a substantial reduction in Lhx2 footprints in both E14 and P2 *Lhx2* cKO samples. The average read coverage was reduced 4-fold in E14 *Lhx2* cKO retinas (Fig. 4A), while in P2 *Lhx2* cKO retinas a 1.5-fold reduction in the average read coverage was observed (Fig. 4B). The reduction in footprint inflection depth was even more pronounced, with an 8-fold reduction observed at E14 and a 1.6-fold reduction at P2. Finally, 96% of all Lhx2 footprints identified in ATAC-Seq profiles of E14 RPCs were lost in E14 *Lhx2* cKO retinas, while 39% were lost at P2 (Auxiliary table 8), and mean footprint signal was significantly reduced at both ages (Auxiliary table 9). This difference between E14 and P2 may partially reflect perdurance of Lhx2 expression at P2 following Cre-mediated deletion from P0 onward. Local accessibility of Lhx2 target sites correlated with Lhx2 occupancy, both at E14 (Fig. S8H) and P2 (Fig. S8I), and significant reductions were observed in *Lhx2* cKO retinas at both ages. Because local accessibility of Lhx2 targets was reduced at both E14 and P2 (Fig. 4C,D upper panel, Fig. S8F,G; Auxiliary table 10), we hypothesized that Lhx2 might play a central role in maintenance of Lhx2-targeted OCRs in RPCs. 40% of all OCRs associated with Lhx2 ChIP-Seq peaks are lost in E14 *Lhx2* cKO samples, while 8% are lost at P2 (Auxiliary table 10). The effect of *Lhx2* loss of function on chromatin accessibility was not merely focal, but extended genome-wide. Overall, 50% of OCRs in the E14 mutants and 22% in P2 mutants were lost (Fig. 4C,D lower panel, Auxiliary table 11), suggesting that Lhx2 is broadly required in RPCs to maintain accessibility of these regions.

### Lhx2 affects local and global chromatin accessibility in RPCs by regulating expression and DNA binding of pioneer TFs

This finding raised the possibility that loss of function of *Lhx2* may disrupt the transcription and/or function of TFs with pioneering activity that are expressed in RPCs, and that these TFs may in turn regulate accessibility of these OCRs. Consistent with this hypothesis, multiple RPC-expressed TFs with predicted pioneering function showed reduced expression and/or reduced accessibility in nearby OCRs in *Lhx2* cKO samples (Fig.4E,F, Auxiliary table 7). Interestingly, a small number of such TFs, such as Dlx2 and Atoh8, showed both reduced accessibility at nearby OCRs and upregulated expression in P2 *Lhx2* cKO samples, implying that these OCRs might mediate Lhx2-depedent transcriptional repression of these genes (Fig. 4F, Auxiliary table 12).

To identify RPC-expressed TFs that might target developmentally dynamic OCRs in parallel with Lhx2, we next analyzed the sequence of OCRs that were selectively active at either E14 or P2. We identified multiple TFs predicted to target RPC-associated OCRs. Some had temporally dynamic expression in RPCs, had known or putative pioneer function (Auxiliary table 13) and were predicted to target Lhx2 ChIP-Seq peaks (Fig. 3C,D)(12).

Overall, TFs that showed enriched expression in E14 or P2 RPCs showed higher predicted pioneering function than TFs that showed similar expression at both ages (Fig. S9). TFs with known or predicted pioneering activity that are enriched in E14 RPCs include *Meis1*. In P2 RPCs, these include members of several TF subfamilies, including SoxE *(Sox8/9)*, NFI *(Nfia/Nfib/Nfix) *(13)*, Ascl1, Hes5*, and KLF *(Klf9/13)* (*14*). Several of these TFs control development of late-born cell types (*8, 15–17*). However, other TFs that showed equivalent levels of mRNA expression in E14 and P2 RPCs – such as Foxm1, Olig2, and Pax6 – preferentially targeted OCRs that were selectively active in either early or late-stage RPCs (Fig. 3C,D; Auxiliary table 8). Multiple TFs expressed in RPCs – such as *Sox2 *(18)**, Smad3, ZFX (Zfhx4) and Etv5 – that exhibit a high pioneer potential were directly transcriptionally targeted by Lhx2 (Auxiliary table 14). These TFs also showed reductions in both footprint count and mean footprint signal (Fig. 4G, Auxiliary table 8, 9) following loss of *Lhx2* function.

### Dynamic Lhx2-dependent regulation of DNA binding by RPC-expressed TFs with predicted pioneer activity

We next conducted a more detailed analysis of our ATAC-Seq data to investigate how loss of function of *Lhx2* at both E14 and P2 affected binding by selected candidate pioneer TFs. As expected, the average insertion frequency at Lhx2 motifs was substantially reduced in both E14 and P2 Lhx2-deficient retinas (Fig. 5A). In line with the mRNA expression patterns of their associated TFs, NF-I and KLF footprints are overrepresented in P2 compared to E14 OCRs, while Sox2 showed comparable occupancy in early and late RPCs (Auxiliary table 8,9). The total number of footprints and the mean signal associated with footprints of gliogenic factors, such as NF-I (Nfia/b/x) were reduced in *Lhx2* cKO, consistent with the disruption of gliogenesis that occurs following *Lhx2* loss of function *(9)*, as were most KLF-related footprints (Fig. 4G, Auxiliary table 8,9).

NF-I, Sox2, KLF, Ascl1 and Hes5 showed a significant reduction in the mean footprint scores at the center of their respective motifs (Fig. 5A-F). For example, 84.4% of the NF-I footprints were lost in the E14 *Lhx2* mutant retinas, as were 93% of the Sox2 associated footprints and 87.6 % of the KLF-related footprints (Auxiliary table 11). The mean footprint signal was also reduced in both E14 and P2 cKO samples (Auxiliary table 12). Among motifs that co-occur with Lhx2, a subset are recognized by TFs with known or predicted pioneer function and show differential expression in early and late RPCs, potentially priming closed chromatin for subsequent opening (Auxiliary table 14).

To better understand the interplay between chromatin organization and Lhx2-dependent regulation of TF binding, we analyzed the nucleosome occupancy around the TF binding sites. Nucleosome occupancy at the center of these motifs was altered in Lhx2 cKO samples, suggesting that Lhx2 may facilitate or stabilize binding by these TFs (Fig. 5G-L). Lhx2 competed for nucleosome occupancy in a manner like that previously described for the non-canonical pioneer factor Nfib *(13)*, with competition was more pronounced at E14 than at P2 (Fig. 5G). In E14 and P2 *Lhx2* cKO retinas, an overall increase in nucleosome coverage occurs, consistent with the reduction in chromatin accessibility observed upon *Lhx2*loss of function. An increase in chromatin compaction was also observed around the Sox2 motif center, although no difference in competition could be detected in *Lhx2* cKO samples (Fig. 5H), suggesting that Sox2 is preferentially recruited to nucleosome-free regions. NFI and KLF factors, as well as Hes5, competed directly for nucleosome occupancy, although no difference was seen on motif centers for NFI and KLF in *Lhx2* cKO samples (Fig. 5I-K). A reduction in chromatin compaction associated with these NFI and KLF sites was seen in the E14 *Lhx2* cKO, while the opposite effect was observed in the P2 *Lhx2* cKO, implying that NFI and KLF factors mediate chromatin unfolding in late-stage RPCs. Finally, nucleosome occupancy by Ascl1 was reduced in the *Lhx2* cKO retinas at both time points, with an effect on flanking regions also observed at P2 (Fig. 5L).

## Discussion

In this study, we report a central role for Lhx2 in the global control of chromatin accessibility in RPCs and identify possible mechanisms by which Lhx2 may regulate gene expression over the course of retinal neurogenesis. First, an unbiased search for oligonucleotides enriched in OCRs found in both early and late RPCs allowed the identification of Lhx2 as the most overrepresented canonical transcription factor. The Lhx2 *cis*-regulatory repertoire changes over the course of neurogenesis, where the selective occupancy of retinal targets is reflected into gene variations upon Lhx2 loss of function, with preferential targeting of genes specifically expressed in early retinal progenitors at embryonic stages, late progenitors and postnatal stages and a lower representation of Muller glia and anterodorsal hypothalamus related genes sets. Changes in gene expression and chromatin accessibility seen following loss of *Lhx2* function are more modest in P2 RPCs than in E14 RPCs. While this fact may partially reflect more extensive perdurance of Lhx2 expression and footprinting that is seen in at P2 RPCs (Fig. 4G), it may indicate that a more diverse, and partially *Lhx2*-independent, transcriptional regulatory network regulates the activity and accessibility of these sequences in late-stage RPCs.

Second, Lhx2 affects local and global chromatin accessibility in RPCs by competition for nucleosome occupancy and through nucleosome remodeling. Lhx2 target sites are often coupled with active enhancer marks, or flanked by active promoter regions within modules of accessible chromatin. *Lhx2* loss of function affects expression of genes involved in the cell cycle checkpoints, DNA replication, axonal dystrophy and the Notch signaling pathway, consistent with previous findings *(7, 9)*.

Third, Lhx2-dependent regulation of both expression and function of pioneer TFs in RPCs appears to contribute substantially to these changes in chromatin accessibility. The regulation of known pioneer factors by Lhx2 is achieved both transcriptionally, as is the case of Lhx2-dependent regulation of OCRs associated with putative cis-regulatory elements associated with the *Sox2* locus, and by stabilizing the binding these TFs to their target sites, as in the case of Sox9, Ascl1 and Hes5. A number of Lhx2-regulated predicted pioneer factors show substantially elevated expression in late-stage RPCs. These include *Hes5, Sox8/9*, and *Nfia*, which play important roles in the control of retinal gliogenesis, a process that is dependent on *Lhx2 *(9)*. Ascl1*, which is also selectively expressed in late-stage RPCs, plays an essential role in conferring neurogenic competence on late-stage RPCs *(15)*. In contrast, other direct Lhx2 targets such as *Pax6* and *Sox2*, like Lhx2, both show broad temporal expression and perform different functions in early and late-stage RPCs *(19–22)*. Some of the known and candidate pioneer factors that are selectively expressed in late-stage RPCs – such as *Nfia/b/x, Sox8/9*, and *Klf4/9/13* -- may preserve chromatin in a highly compacted state at embryonic states, while mediating chromatin unfolding later in development.

The precise molecular mechanism by which Lhx2 mediates these developmentally dynamic changes in chromatin accessibility is unclear. However, recent studies of Lhx2 function in developing cortex have identified multiple histone modifying enzymes that interact with Lhx2 *(23)*. Interestingly, these include NuRD complex components that are associated with transcriptional repression and reduced chromatin accessibility, such as LSD1, HDAC2, and RBBP4. In RPCs, Lhx2 may instead recruit different sets of histone remodeling enzymes that promote chromatin accessibility, which should be readily identifiable through biochemical analysis.

## Figure legends

**Fig 1. Pairwise comparison of ATAC-Seq data identifies OCRs from early and late RPCs, with a broad overrepresentation of Lhx2-related motifs. (A)** Workflow adopted for epigenomic profiling of RPC **(B)** Venn diagram represents open chromatin regions identified in early and late-stage RPCs **(C)** Clustering of TF motifs based on similarity (left, Jaspar 2016 non-redundant vertebrates core). The heatmap indicates similarity for each pairwise comparison. The color bars along the axis represent different TF subfamilies. Inset represents a homeobox cluster that was enriched in OCRs in RPCs (scale= 1 node/pixel) **(D)** Relative expression by RNA-Seq for homeobox TFs **(E)** Representative Lhx2-related logos are shown (MA0700.1, k-mer sig=300, e-value=1e-300) **(F)** Binomial scoring of known motifs in chromatin regions accessible in early and late RPCs **(G)** Lhx2 footprints from OCRs identified in early and late-stage RPCs **(H)** Custom tracks of RPC-derived ATAC-Seq profiles and aged-matched Lhx2 ChIP-Seq feature the *Vsx2* locus.

**Figure 2. Lhx2 regulates cell cycle genes and the Notch signaling pathway by targeting promoters and non-coding elements in nucleosome-free regions associated with active enhancers (A)** Lhx2 known motif density for E14 and P2 ChIP-Seq peaks. **(B)** Lhx2 peak association with specific genomic regions (log_2_-fold enrichment). **(C-F)** Enrichment for cell-specific gene expression patterns is shown for Lhx2 peaks that are shared between E14 and P2 (C), associated with stage-specific Lhx2 peaks (D), and all Lhx2 peaks detected at E14 **(E)** and P2 (F). RNA-Seq from age-matched Lhx2 cKO retinas identifies Lhx2-dependent genes sets associated with at least one ChIP-Seq peak. Asterisks refer to p-values after Bonferroni-Hochberg correction. **(G,H)** Heatmaps of raw reads from Lhx2 ChIP-Seq peaks are plotted across nucleosome centered regions identified using age-matched ATAC-Seq samples. Each row represents a 3 kb window (1.5 kb each direction) centered at the maximum read pile-up. For Lhx2 motif occurrence in OCRs, refer to Auxiliary table S8. Lhx2 motifs at open chromatin regions co-directionally distribute with RNA-Seq raw reads. Metaprofiles of the class II enhancer-associated H3K27ac marks were compiled at actively transcribed regions from the E14 GFP-positive **(G)** and P2 GFP-positive RPCs **(H)** (replicates in grey scale, background in light grey). **(I,J)** Binomial scoring of Gene Ontology (GO) functional categories that are enriched in genes associated with OCRs that overlap with Lhx2 peaks at E14 **(I)** or P2 **(J)**.

**Figure 3. Binding sites for transcription factors with predicted pioneer function cooccur with Lhx2 peaks (A,B)** Hierarchical clustering of Lhx2 ChIP-Seq regulatory motifs and assigned representative logos are represented at E14 **(A)** and P2 **(B)** (linkage=average; similarity threshold cor =0.6, ncor=0.4, w=5). The most enriched cluster comprises Lhx2 and multiple close variations of the Lhx2 motif. **(C,D)** Known TF motifs found preferentially enriched in either E14 or P2 Lhx2 ChIP-Seq peaks, and identified by pairwise comparison, are shown. The percentages of Lhx2 ChIP-Seq peaks that contain the indicated motif are listed, along with the background frequency for the motif in question. The relative expression level of the corresponding TF mRNA at E14 or P2 is indicated by the red/blue color gradient.

**Figure 4. Lhx2 affects local and global chromatin accessibility in RPCs by regulating expression and DNA binding of pioneer TFs (A,B)** ATAC-Seq read distribution plotted relative to the center of Lhx2 footprints, and average inflection depth for these footprints in both controls and *Lhx2* mutants. **(C,D)** Heat maps of raw reads from Lhx2 ChIP-Seq and age-matched OCRs, identified by ATAC-Seq of purified RPCs, are plotted in a 3kb window relative to the center of OCRs from control and *Lhx2* cKO retinas, with the difference in signal between these two conditions shown at right. **(E,F)** Paired variation of RNA-Seq signal and local chromatin accessibility in E14 **(E)** and P2 **(F)** control and *Lhx2* cKO samples are shown for loci encoding TFs that are regulated by Lhx2. Individual representative TFs are highlighted by arrows. **(G)** Footprint counts for individual TFs with predicted pioneer activity that are associated with Lhx2 target sites are shown in both control and *Lhx2* mutants from E14 and P2 retina.

**Fig 5. Metaprofiles of footprinting and nucleosome occupancy data reveal developmentally dynamic regulation by Lhx2 of DNA binding by RPC-expressed TFs with predicted pioneer activity. (A-F)** ATAC-Seq average cut profiles feature footprints for Lhx2, Sox2, NF-I (Nfia/b/x), Klf4/9, Hes5 and Ascl1 from both E14 and P2 control and *Lhx2* cKO samples. **(G-L)** Nucleosome occupancy at the center of each motif is reported for Lhx2, Sox2, NF-I (Nfia/b/x), Klf4/9, Hes5, and Ascl1 in both E14 and P2 control and *Lhx2* cKO samples.

## Materials and Methods

### Experimental design

The experimental pipeline involves the generation and integration of high-throughput sequencing libraries from purified fractions of murine retinal progenitors cells (RPCs). Fluorescence-activated cell sorting (FACS) was adopted to isolate RPCs from mice expressing the RPC-specific *Chx10-Cre:eGFP* transgene (*11*) from embryonic day (E)14 and postnatal day (P)2 retina, representing early and late stages of RPC competence, respectively. Flow-sorted cells were profiled by RNA-Seq and ATAC-Seq, relying on direct *in vitro* transposition of sequencing adapters into native chromatin. ChIP-Seq was then performed on a select transcription factor candidate, identified in the preliminary screening of the ATAC-Seq derived open chromatin regions. ChIP-Seq binding sites for the candidate factor were ultimately integrated and compared with the age-matched RNA-Seq and ATAC-Seq profiles from control and *Lhx2* cKO retinas. For each experimental condition involving a point-source factor and broad regions (ChIP-Seq and ATAC-Seq, respectively) a minimum of 20 Millions uniquely mappable reads or ≥10 millions uniquely mappable reads for each biological replicate were collected, according to the ENCODE’s guidelines. For point-source datasets, non-redundant mapped reads were retained for downstream analysis.

### Animals

CD1 mice of either sex were euthanized at embryonic day 14 (E14) and postnatal day 2 (P2) according to Johns Hopkins IACUC-approved protocols. Timed pregnant CD-1 mice were obtained from Charles River Laboratories. *Chx10-Cre:GFP* mice (*11*) were purchased from the Jackson Laboratories. Retinas were freshly dissected, incubated in a suspension of papain and DNAse for 30 min at 37 C, inactivated with bovine serum albumin, resuspended in equilibrated Earle’s balanced salt solution and subject to fluorescence activated cell sorting (FACS) to 98-99% purity, with viability assessed by propidium iodide exclusion. Cell fractions were collected on poly-D-lysine coated slides, fixed in 4% paraformaldehyde for 10 min, permeabilized in TritonX-100 and stained for Chx10 (Cat.# X1179P, Exalpha), GFP (Cat.# 600-101-215, Rockland), Ki67 (Cat.# RM-9106-S1,Thermo Scientific) or Ccnd1 (Cat.# SC-450, Santa Cruz). The brightest fraction, which consistently showed 4-fold higher mean intensity for GFP relative to the dim fraction, was always retained for subsequent processing and hereafter referred to as GFP-positive, RPC-enriched fraction. RNA-Seq analysis revealed a substantial enrichment of RPC-specific markers in the GFP-positive fraction relative to the age-matched GFP-negative fraction. Examples of this include (number indicates level in GFP-positive relative to age-matched GFP-negative cells): Ccnd1, E14=22.9, P2=2.9; Mki67, E14=19.7, P2=3.1; Vsx2, E14=47.7, P2=2.8; Fgf15, E14=45.4, P2=3.4.

*Lhx2* conditional embryonic knockouts were obtained by crossing *Chx10-Cre:GFP* with *Lhx^lox/lox^*mice, and harvesting at E14 *(24)*. Postnatal *Lhx2* knockouts were generated by electroporation of pCAG-Cre-GFP construct into P0.5 wild type CD1 animals or *Lhx2^lox/lox^* animals. Retinas were harvested at P2, dissociated, and GFP-positive electroporated cells were isolated by FACS. Overall electroporation efficiency was 2-3%.

### ATAC-Seq, RNA-Seq, and ChlP-Seq analysis

Chromatin derived from flow-sorted *Chx10-CreGFP+ve* and GFP-ve retinal fractions was processed as previously described *(25)*. Briefly, chromatin was extracted and processed for Tn5 mediated tagmentation and adapter incorporation, according to the Manufacturer’s protocol (Nextera DNA sample preparation kit, Illumina®) at 37 C for 30 min. Reduced-cycle amplification was carried out in presence of compatibly indexed sequencing adapters. The quality of the libraries was assessed by fluorometric DNA incorporation based assay (Thermo Fisher Scientific™) and automated capillary electrophoresis (Agilent Technologies, Inc.) and up to 4 samples per lane were pooled and run as 50 bp paired ends on a HiSeq2500 Illumina sequencer.

RNA was processed with Qiagen RNAeasy Mini kit, subject to DNAse digestion, and samples with a minimum RNA integrity number (RIN) of 7 were further processed for sequencing. Libraries were prepared using Illumina TruSeq RNA Sample kit (Illumina, San Diego, CA) following manufacturer’s recommended procedure. Briefly, total RNA was denatured at 65°C for 5 minutes, cooled on ice, purified and incubated at 80°C for two minutes. The eluted mRNA was fragmented at 94°C for 8 min and converted to double stranded cDNA, end repaired, A-tailed, and ligated with indexed adaptors and run on a MiSeq Illumina sequencer. The quality of the libraries was assessed by fluorometric RNA incorporation based assay (Thermo Fisher Scientific™) and automated capillary electrophoresis (Agilent Technologies, Inc).

ChIP was performed as described previously *(26)*. Whole dissected retinas were dissociated in a collagenase I suspension, cross-linked in 1% formaldehyde for 15 min, and quenched in 125 mM glycine. The extracted nuclei were sheared to produce 100–500 bp fragments by means of probe sonication. Chromatin was immunoprecipitated with anti-Lhx2 (Cat.# SC-19344, Santa Cruz Biotechnology), rabbit anti-H3K27Ac ((Cat.# ab4729, Abcam), or the corresponding isotype controls (Abcam); retained on agarose beads (Invitrogen), washed and purified by organic extraction. Success of both anti-Lhx2 and anti-H3K27Ac ChIP was confirmed using SYBR qRT-PCR (Agilent Technologies) that performed using previously described primer pairs that amplify both cis-regulatory regions that exhibit Lhx2-binding sites and syntenic unbound regions *(9)*. Libraries were processed according to the manufacturer’s protocol (TruSeq Nano DNA Library Prep Kit). The quality of the libraries was assessed by fluorometric DNA incorporation based assay (Thermo-Fisher) and automated capillary electrophoresis (Agilent Technologies) and libraries (100-150bp single read, paired ends) were run on a HiSeq2500 Illumina sequencer.

### Peak calling and retinal gene ontology analysis

RNA-Seq reads were aligned to the mouse transcriptome (mm9 UCSC build) using Tophat2 *(27)*, and differentially expressed genes (DEGs) were identified by Cuffdiffs *(28)*, imposing a cutoff q-val = 0.05 for pairwise comparison.

Bowtie2 was used for ChIP-Seq and ATAC-Seq reads alignment on the mouse genome (mm9) *(29)*. Uniquely mappable reads from ChIP-Seq were retained for peak calling by MACSs *(30)* (band width = 300, mfold =5, 50, d = 200, max tags per position = 1, min FDR q-val cutoff = 1E-02, λ = 1000-10000 bp).

Open chromatin regions were identified in ATAC-Seq data using MACS2 *(30)*. Correlations between open chromatin states was identified by pairwise comparison of normalized reads counts in E14 and P2 GFP-positive flow-sorted retinal progenitor cells was done with Jaccard *(31)*. The Jaccard index was estimated as the number of peaks that overlap between two peak files, divided by the union of the two files. Footprints and nucleosomes were identified as described previously *(12, 32, 33)*. Annotation was done by proximity to the closest transcription start site.

High-confidence ChIP-Seq peaks were identified from at least 2 experimental replicates (Poisson p-val threshold = 0.0001, min FE=4, FDR=0.001, max tags per position =1, normalization to input or isotype control) and subject to comparison with ATAC-Seq peaks (hypergeometric test, ln p-val) where co-occurrence was defined by physical overlap, allowing a max distance of 20 bp from peak summits over 3000 bp for confocality. For direct comparison of ChIP-Seq and ATAC-Seq data, high confidence peaks with the highest differential in accessibility between nucleosomal units and flanking nucleosomes-free, transposon-accessible regions, and a minimum distance of 300bp were identified from two ATAC-Seq replicates.

Genes that showed enriched or specific expression in retinal progenitors were identified generated based on the RNA-Seq data generated in this study (for RPC-specific expression). Genes that showed cell-type enriched or specific expression in adult retina were extracted from other published studies using either RNA-Seq or microarray-based analysis of isolated cells, or *in situ* hybridization or immunohistochemistry of intact retina. Association of ChIP-Seq peaks with genes expressed in specific retinal cell types was evaluated by Fisher’s exact test and corrected for multiple hypothesis testing (Bonferroni).

### Motif enrichment analysis

Hierarchical clustering of probabilistically assigned motifs (Jaspar 2016 non redundant core vertebrates) *(34)* was done with the following parameters (linkage=average; similarity threshold cor =0.6, ncor=0.4, rl=5, where ncor is Pearson’s correlation (cor) by relative alignment length (rl) divided by the overall alignment length. (e-val= pval * enriched oligos) *(35)*.

For ChIP-Seq analysis, binomial probability analysis of TF motifs was calculated in the overall set of high confidence peaks (minimum of 2 normalized replicates). The regions size was empirically determined and motifs were found by cumulative binomial distribution of known position weight matrices assuming a random representation of decamers. Motif finding was performed on the repeat-masked mm9 murine genome, optimized for the top enriched 20 putative motifs, randomized and repeated twice to estimate FDR *(36)*.

### Footprint analysis

FIMO *(37)* was used to scan the genome for candidate binding sites of different transcription factors. We then used BPAC to identify the actual binding sites among these candidate sites *(33)*. The number of active binding sites was analyzed at both E14 and P2. Genome-wide changes in footprint counts and nucleosome occupancy for individual transcription factor motifs were estimated for all candidate TF binding sites at both E14 and P2. Nucleosome occupancy was estimated using NucleoATAC *(32)*. Footprints scores were calculated for Lhx2, Klf9/13, NF-I (Nfia/b/x), and Sox2. T-test on mean footprints scores distributed 200bp within of motif site centers were calculated for the paired control and *Lhx2* cKO conditions.

### Statistical analysis

Co-occurrence statistics for point-source and broad regions of interest was computed by hypergeometric test with a default minimum overlap of 1 bp, unless otherwise specified. Coverage was adopted as reproducibility metrics for ChIP-Seq and ATAC-Seq experimental replicates (fraction reads/10^7^ non-redundant uniquely mappable reads) and FPKM for RNA-Seq, where correlation was reported by Pearson’s or Spearman’s coefficient. Binomial probability analysis of regulatory transcription factors motifs was applied genome wide to identify enriched position weight matrices (PWMs) and clustered by average linkage. Fisher’s exact test was adopted to compute gene ontology enrichment and corrected for multiple hypothesis (Bonferroni-Hochberg) and RNA-Seq derived gene sets from flow sorted retinal cell fractions and control versus experimental conditions *(Lhx2* cKO) with q-val (FDR) < 0.05 were retained for downstream analysis, unless otherwise specified. Two-tailed t-test was adopted to compare the average footprint counts for candidate pioneer factors in control and *Lhx2* cKO samples.

## Acknowledgments

**General**: We thank Hongkai Ji for help with statistical analysis, Anand Venkataraman and Wendy Yap for comments on the manuscript, Hong Wang for help with dissections, the JHMI Deep Sequencing and Microarray core, the JHU GRCF High Throughput Sequencing facility, and the JHSPH Flow Cytometry and Cell Sorting Core Facility.

## Author contributions

C.Z. carried out experiments and data analysis. S.L, J.W. and J.Q carried out analysis. C.Z., J.Q. and S.B. conceived the study and wrote the paper.

## Data and materials availability

Auxiliary data and files have been deposited in GEO, Accession Number GSE99818. All data needed to evaluate the conclusions in the paper are present in the paper and/or the Supplementary Materials. Additional data is available from the authors upon request.

